# Pulse of α-2-macroglobulin and lipocalin-1 in the pregnant uterus of European polecats (*Mustela putorius*) at the time of implantation

**DOI:** 10.1101/088609

**Authors:** Heli Lindeberg, Richard J.S. Burchmore, Malcolm W. Kennedy

**Affiliations:** Natural Resources Institute Finland (Luke), Green Technology, Halolantie 31 A, FIN-71750 Maaninka, Finland.; Institute of Infection, Immunity and Inflammation, and Glasgow Polyomics, College of Medical, Veterinary and Life Sciences, University of Glasgow, Garscube Campus, Glasgow G61 1QH, Scotland, UK.; Institute of Biodiversity, Animal Health and Comparative Medicine, and the Institute of Molecular, Cell and Systems Biology, Graham Kerr Building, College of Medical, Veterinary and Life Sciences, University of Glasgow, Glasgow G12 8QQ, Scotland, UK

**Keywords:** European polecat, ferret, *Mustela putorius*, uterine secretions, pregnancy, implantation, proteomics

## Abstract

Uterine secretory proteins protect the uterus and conceptuses against infection, facilitate implantation, control cellular damage resulting from implantation, and supply embryos with nutrients. The early conceptus of the European polecat (*Mustela putorius*) grows and develops free in the uterus until implanting at about 12 days after mating. Using a proteomics approach we found that the proteins appearing in the uterus leading up to and including the time of implantation changed dramatically with time. Several of the proteins identified have been found in pregnant uteri of other placental mammals, such as α_1_-antitrypsin, serum albumin, lactoferrin, cathepsin L1, uteroferrin, and ectonucleotide pyrophosphatase. The broad-spectrum proteinase inhibitor α_2_-macroglobulin rose from relatively low abundance initially to dominate the protein profile by the time of implantation. Its functions may be to limit damage caused by the release of proteinases during implantation, and to control other processes around the site of implantation. Lipocalin-1 (also known as tear lipocalin) has not previously been recorded as a uterine secretion in pregnancy, and also increased substantially in concentration. If polecat lipocalin-1 has similar biochemical properties to the human form, then it may have a combined function in transporting or scavenging lipids, and antimicrobial activities. The changes in the uterine secretory proteome of Euroepan polecats may be similar in those species of mustelid that engage in embryonic diapause, but possibly only following reactivation of the embryo.

## 1. Introduction

A conceptus may implant soon after its arrival in the uterus (as in humans and muroid rodents), or may remain free in the uterus for several weeks before implanting (as in equids) [1]. Intermediate to these extremes are the conceptuses of many Carnivora and Artiodactyla which implant 10-20 days after mating. The delay is attributed to time required for mating-induced ovulation, travel through the fallopian tubes, and a variable period during which the blastocyst develops without close cellular contact with maternal tissues [1]. In some species of mustelid, ursids, phocids, and roe deer, implantation is postponed for several months, during which time the conceptus enters obligatory diapause at the blastocyst stage and must be maintained and protected until reactivation is permitted by the mother [2-5]. The European polecat, *Mustela putorius* [6], fits into the intermediate group, with an embryonic development period (without embryonic diapause) of approximately 12 days preceding implantation.

Proteins secreted into the non-pregnant uteri of eutherian mammals have a range of presumptive functions such as maintenance of the mucocutaneous surface of the endometrium, antimicrobial protection, receptivity to sperm and the subsequent conceptus, and nutrition of an embryo. Distinct differences have been found among mammal groups in the proteins secreted into the uterus before placentation, although there are commonalities [7-12]. These proteins have been divided mainly into those involved in nutritional support for the conceptus, or in facilitating and controlling implantation events. Perhaps the most notable example of the nutritional function is the uterocalin/P19 protein in horses. This protein appears to deliver essential lipids such as retinol and polyunsaturated fatty acids across the glycoprotein capsule to the equine conceptus [13-15]. Lipids present a problem in their transfer from mother to conceptus because they tend to be insoluble, and in some cases susceptible to oxidation damage unless protected within a protein binding site. Uterocalin is also notable in that its primary structure is enriched in essential amino acids, which doubles its function as a nutrient source for the developing embryo [15]. Other examples of lipid carriers in uterine secretions include a modified form of plasma retinol binding protein, and serum albumin which binds a range of small molecules, fatty acids in particular [11, 16, 17]. Of those proteins considered to influence implantation events, the broad-spectrum proteinase inhibitor α-2-macroglobulin (α_2_M) is secreted into the pregnant uterus around the time of implantation in several species, and, among other roles, is thought to limit tissue damage during implantation and to control local inflammatory responses [7-12, 18, 19].

Overall, there is considerable diversity in the suite of proteins secreted into pre-placentation uteri by different clades of mammals, as exemplified by equids, artiodactyles and Carnivora, all of which exhibit distinctive repertoires [7-10, 12]. We investigated the proteins found in uterine secretions of pregnant European polecats in order to explore the preparatory events that lead up to, and at, implantation. Analysis of chronological samples indicated that progressive and dramatic changes in the protein profile occur as the time at which implantation would occur approaches. Some of the proteins have been found in pre-implantation uterine flushes of other species (notable among which is α-2-macroglobulin), although not in the same combination or relative concentrations. At least one protein, lipocalin-1, which we found to peak in abundance at implantation, has previously not been reported as a uterine secretory protein. Not only does this study demonstrate that mustelids exhibit a different but overlapping repertoire of implantation-related uterine secretions from other species groups, but it may also provide a general framework for investigation of what happens during embryonic quiescence and subsequent reactivation in species that engage in embryonic diapause.

## 2. Materials and Methods

### 2.1 Animals and sample collection

The subjects used in this study descended from a population of 70 wild European polecats (*Mustela putorius*) captured in the 1970s and interbred with captive 200 imported domestic ferrets (subspecies *Mustela putorius furo*). The resulting population forms the basis for all polecats farmed for the fur industry and research in Finland. The animals were maintained at the Kannus Research Farm Luova Ltd., in Kannus, Finland. They were held in individual cages measuring 70 x 30 x 38 cm (length x width x height) with nest boxes measuring 40 x 29 x 32 cm which were accessible to the females but external to the main cage areas. During the breeding season, the animals were exposed to outdoor temperatures and light conditions: mean 2°C and 15 hours light in April, 9°C and 16 hours light in May, 14°C, and 20 hours light in June. The animals were fed according to standards for breeding farmed polecats in Finland (prepared to provide at least 1150-1250 kcal/kg, 40-50% protein, 30-32% fats, and 15-25% carbohydrates, supplemented with vitamins and minerals), and had water provided ad libitum.

The time of oestrus was estimated by the physical appearance of the vulva. Males used for mating were known to have been fertile the previous year, and matings were visually confirmed (a tie observed). Animals were randomly selected for sample collection on days 4, 6, 7, 9, 12 and 14 days after mating, and those sampled on days 4 and 14 were found not to have become pregnant and thereby acted as mated but non-pregnant controls. Each animal was anesthetized with medetomidine (Dorbene^®^; Laboratorios syva s.a., León, Spain) and ketamine (Ketalar^®^; Pfizer Oy, Helsinki, Finland), and the uterus removed under sterile surgical conditions. The animals were subsequently euthanized with T-61 (Intervet International B.V., Boxmeer, the Netherlands) while still under anesthesia. Once the uterus was removed, the tips of each horn were opened, and the cut ends cleared of blood. The cervix was closed with forceps, and the uterus flushed with 5 ml sterile physiological saline through one horn and out the other, with care to avoid blood contamination. The flush fluid was filter-sterilised and frozen at -20 °C in approximately 2 ml aliquots. Pregnancy was confirmed by the presence of developing embryos in uterine flushes.

All of the work was approved by the National Animal Experiment Board (Eläinkoelautakunta, ELLA) of Finland.

### 2.2 Protein electrophoresis

1-dimensional vertical sodium dodecyl sulphate polyacrylamide gel electrophoresis (SDS-PAGE) was carried out using the Invitrogen (Thermo Scientific, Paisley, UK) NuPAGE system using precast 4-12% gradient acrylamide gels, with β-mercaptoethanol (25 μl added to 1 ml sample buffer) as reducing agent when required. Gels were stained for protein using colloidal Coomassie Blue (InstantBlue, Expedion, Harston, UK) and images of gels were recorded using a Kodak imager. Electronic images were modified only for adjustment of contrast and brightness. Pre-stained molecular mass/relative mobility (M_r_) standard proteins were obtained from New England Biolabs, Ipswich, MA, USA (cat. number P7708S). Samples were run under non-reducing and reducing conditions using β2 mercaptoethanol as reducing agent. Selected gel slices were taken from non-reduced, Coomassie Blue-stained gels and processed for application to fresh gels under reducing conditions. Samples were concentrated where required using a Vivaspin 3,000 MWCO PES (Sartorius, Epsom, Surrey, UK) centrifugal concentrator device operated according to the manufacturer’s instructions.

### 2.3. Proteomics

Stained protein bands were excised from preparative 1-dimensional gels and analysed by liquid chromatography-mass spectrometry as previously described [20]. Protein identifications were assigned using the MASCOT search engine to interrogate protein and gene sequences in the NCBI databases and the *Mustela putorius* genome database [21] and linked resources, allowing a mass tolerance of 0.4 Da for both single and tandem mass spectrometry analyses. BLAST searches, or searches of genome databases of other Carnivora (e.g. dog and giant panda), were carried out to check the annotations. Prediction of secretory leader peptides and their cleavage sites were carried out using SignalP software (http://www.cbs.dtu.dk/services/SignalP/;[22]), and molecular masses calculated using ProtParam (http://web.expasy.org/protparam/).

## 3. Results

### 3.1 Sequential changes in the protein profile of pre-implantation uterine secretions

The protein profiles of all the uterine flush samples collected from days 4 to 14 after mating are shown in the SDS-PAGE analysis shown in Supplementary Information Figure S1. The disparities between overall protein concentrations amongst the samples could be due to differences in efficiency of flushing, changes in the total tissue volume, differences between individual animals’ secretion volumes, or changes in secretory activity. In order to improve comparability, selected samples were concentrated as described above and/or the volume of loaded sample adjusted to approximately equalise the intensity of the band at ca. 65 kDa M_r_ (band C; Figure 1), suspected (and subsequently confirmed by mass spectrometry), to be serum albumin from its size and slower migration under reduction.

**Figure 1.**
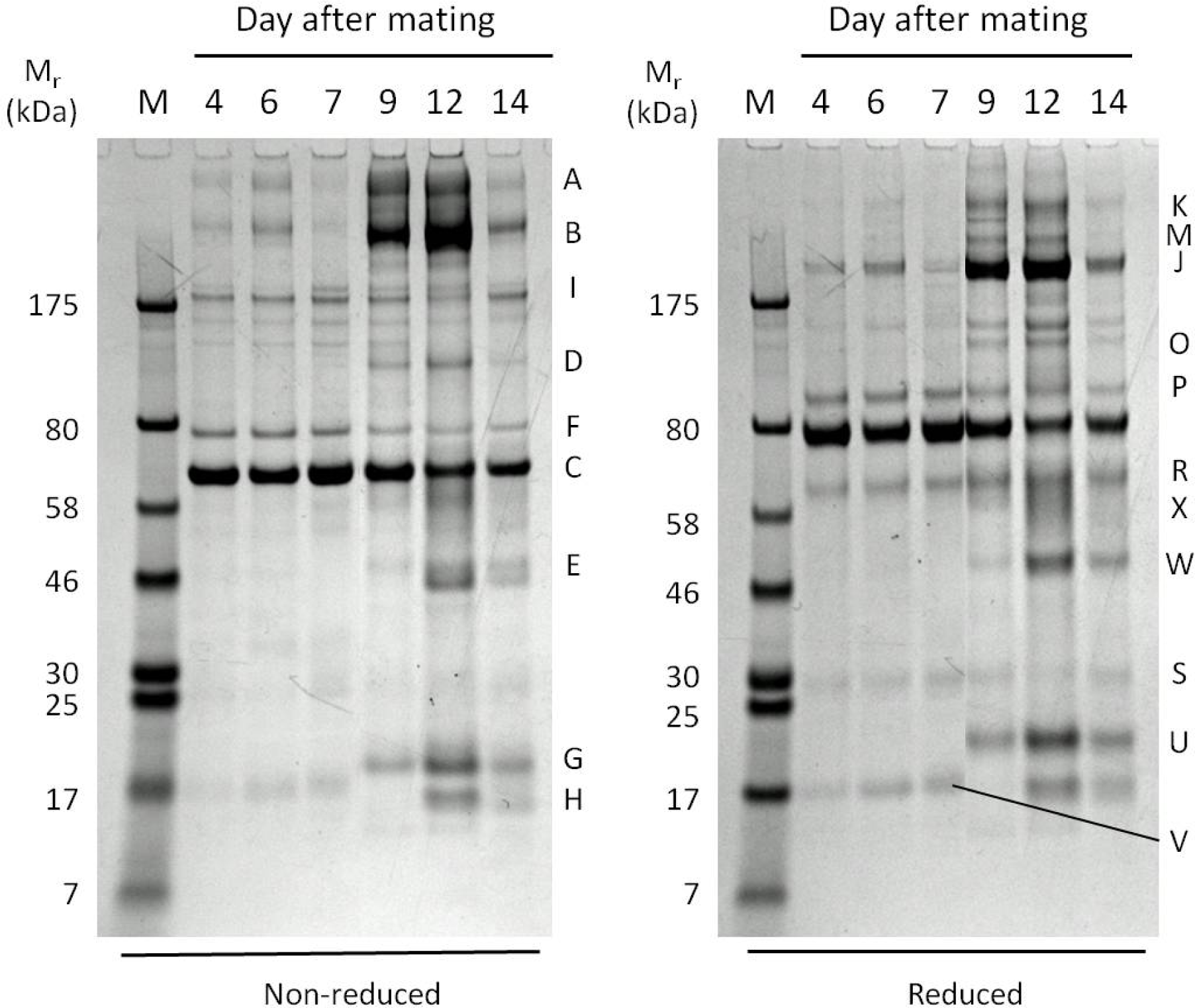
Changes in European polecat uterine secretory proteins with time after mating. Animals sampled on days 4 and 14 were non-pregnant so act as controls. The sample volumes were adjusted to normalize the intensity of the strong band at approximately 65 kDa (serum albumin) among tracks. See Figure S1 for SDS-PAGE of all of the samples collected and upon which the adjustments were based. Gel band codes indicated by letters and are referred to in the text and in Table 1. M – marker/calibration proteins with relative mobilities (M_r_) as indicated in kilodaltons (kDa).

Standardization of protein concentration to serum albumin revealed dramatic changes in the chronological protein profile of pregnant uterine flushes, in particular the proteins in bands A, B, D, E, G, U, and W (Figure 1). The pregnant animals on days 6 and 7 showed few, if any, differences from the mated but non-pregnant animal at day 4, but by day 9 considerable differences were evident between pregnant and non-pregnant individuals.

In order to explore relationships between some of the proteins in the most intense bands, gel slices were excised from a non-reduced gel, and the proteins in them subjected to electrophoresis under reducing conditions. This showed that bands A and B reduced to a similar set of proteins, suggesting that they were related or identical (Figure 2). The sizes of the resulting fragments were not commensurate with post-reduction cleavage products of large multi-chain immunoglobulins (IgM and IgA) which might be expected to appear in uterine secretions. Protein band C, which was used to standardize the loading protein concentrations of samples for Figure 1, migrated more slowly when reduced, which is typical of serum albumin. The change in its migration is usually attributed to cleavage of its intramolecular disulphide bond such that the unfolded protein is retarded in its migration. Similar behaviour was exhibited by band G protein, but the others separated as under non-reducing conditions. The reduction of band C protein uncovered a different protein that was subjected separately to proteomic analysis (band R).

**Figure 2.**
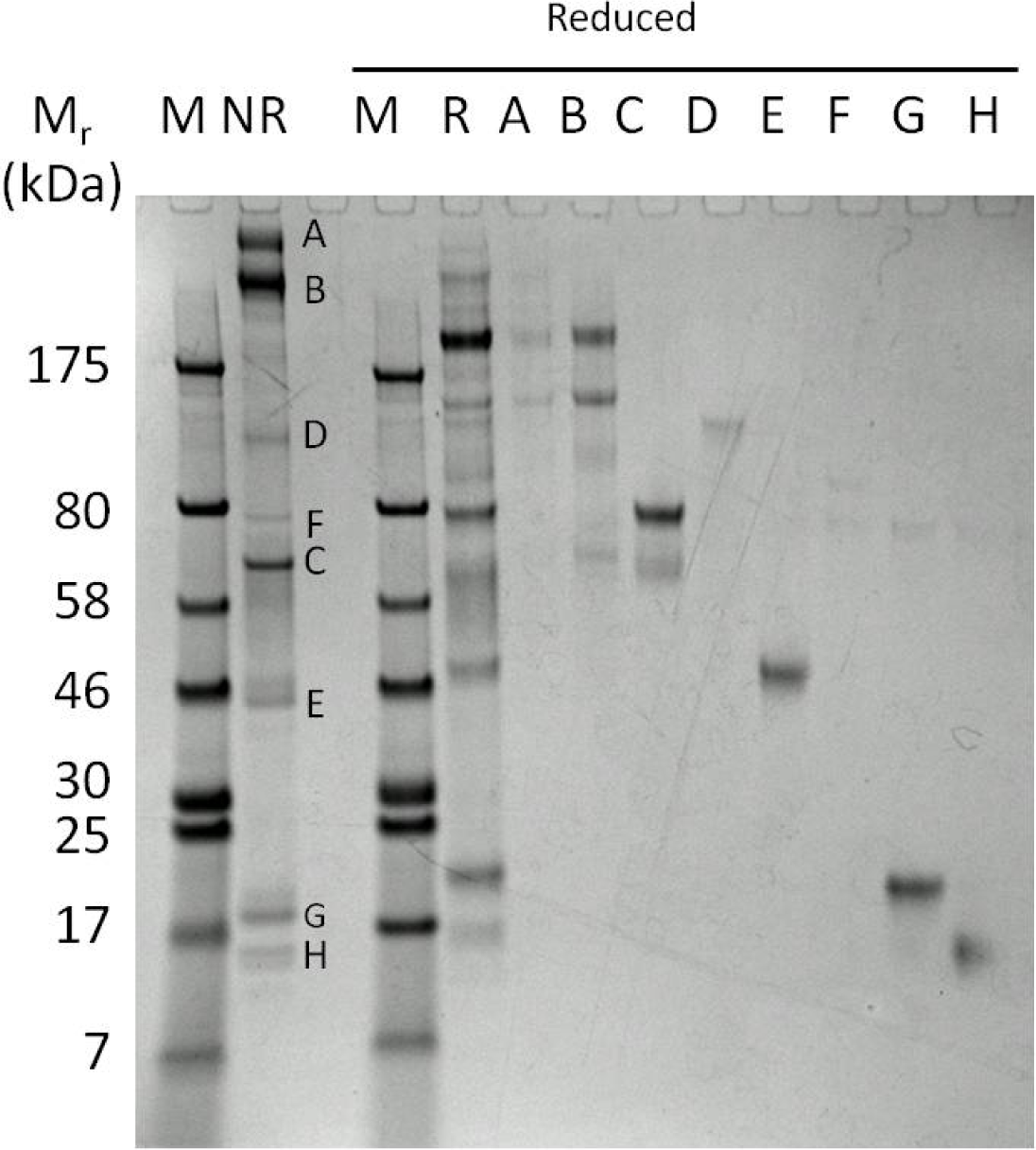
Subunit composition of major proteins in European polecat uterine flush following protein reduction. The labelled bands were excised from a non-reducing preparative SDS-PAGE gel of a uterine flush sample taken from a pregnant animal on day 12 after mating and re-run under reducing conditions. The letter codes for each band are consistent with those in Figure 1. Protein samples were run under reducing or non-reducing (NR) conditions. M – marker/calibration proteins with relative mobilities (M_r_) indicated in kilodaltons (kDa).

### 3.2 Proteomics

Protein bands whose concentrations changed dramatically with time after mating, or whose concentrations remained constant, were selected for proteomic analysis from both reducing and non-reducing gels (Figure 1). The separation between the bands in the SDS-PAGE gels was sufficient to allow proteomic identification of the proteins with high confidence, all directly from the *M. putorius* genome-derived protein sequence database. The deductions were unchanged upon checking by BLAST searching of genomic and other databases for other Carnivora. Table 1 lists the identifications and also the MASCOT scores and peptide matches of the proteins, and illustrates the high quality of the matches.

**Table 1.**
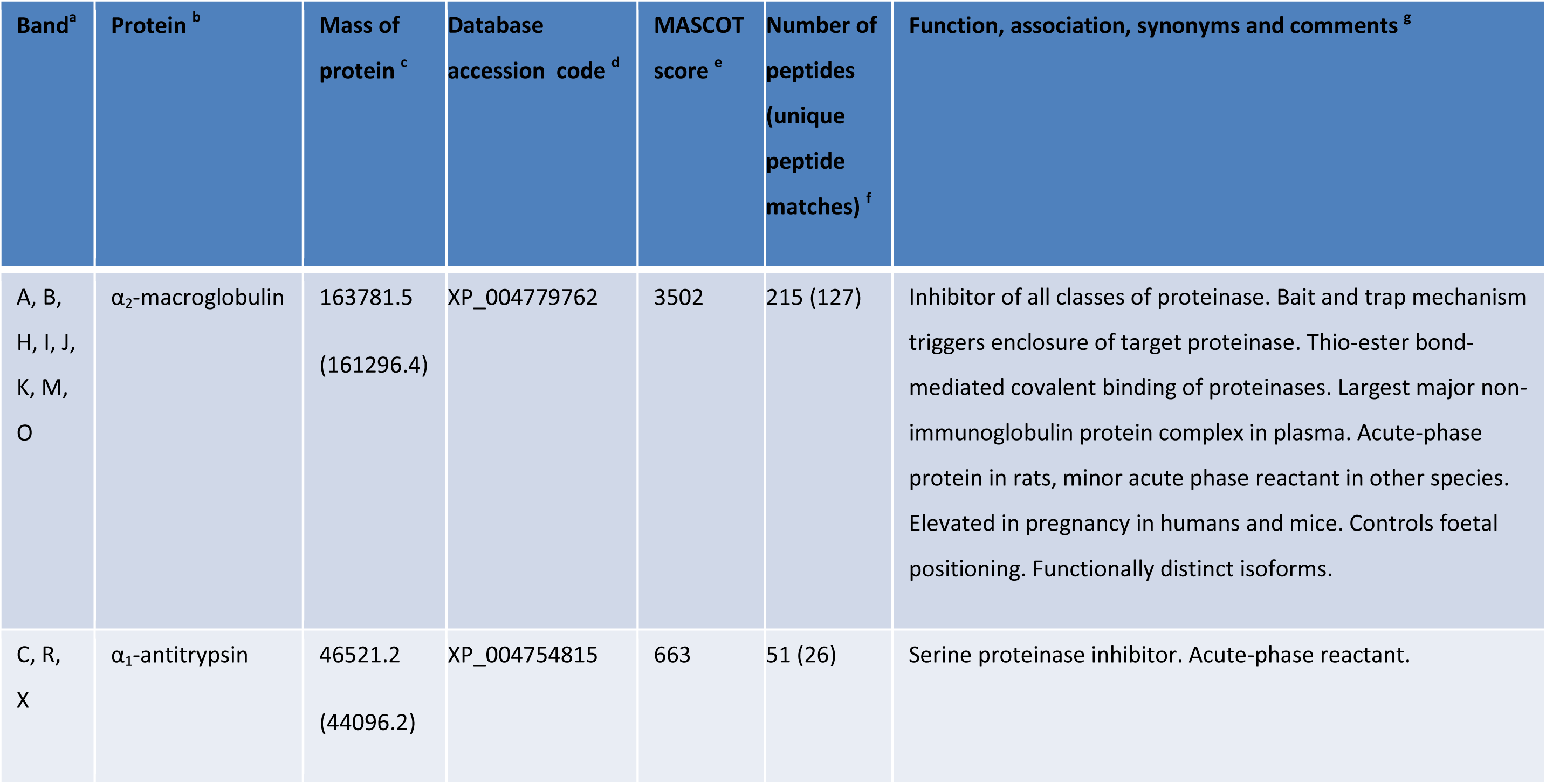
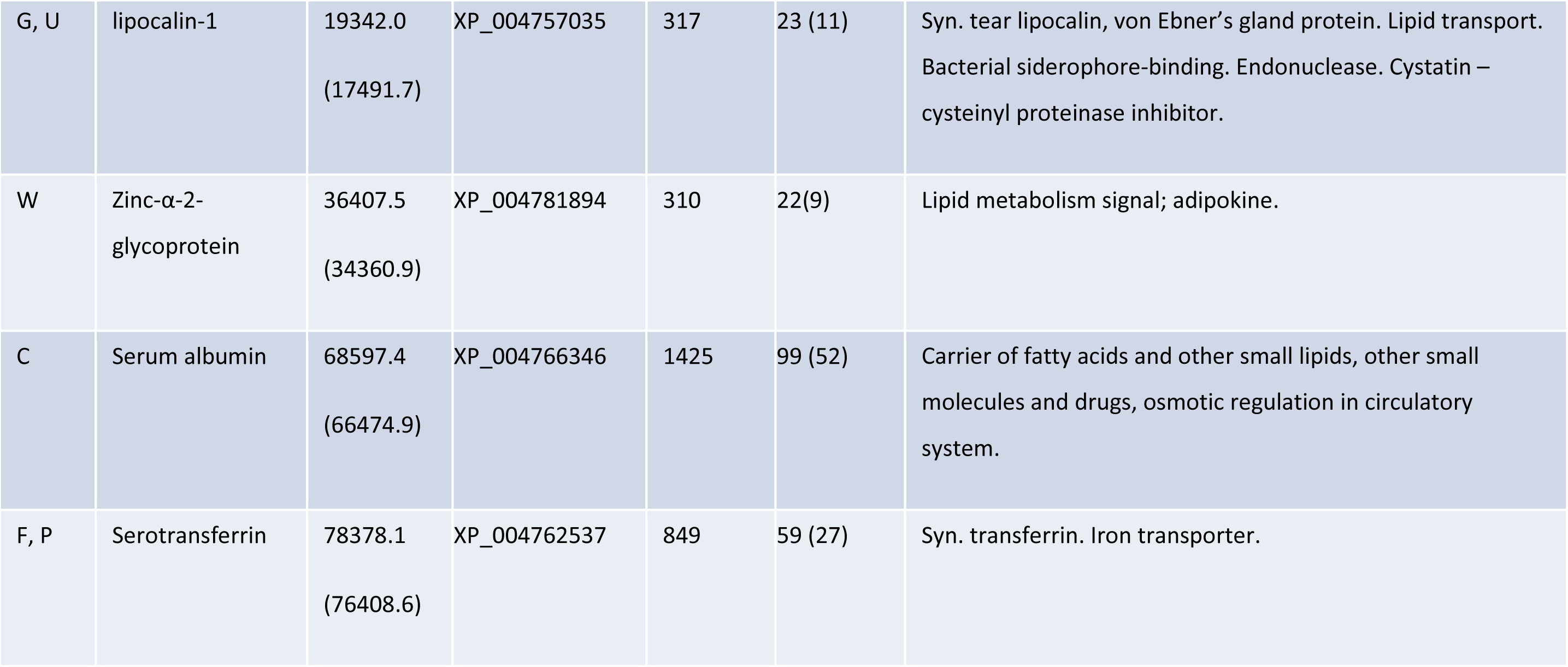
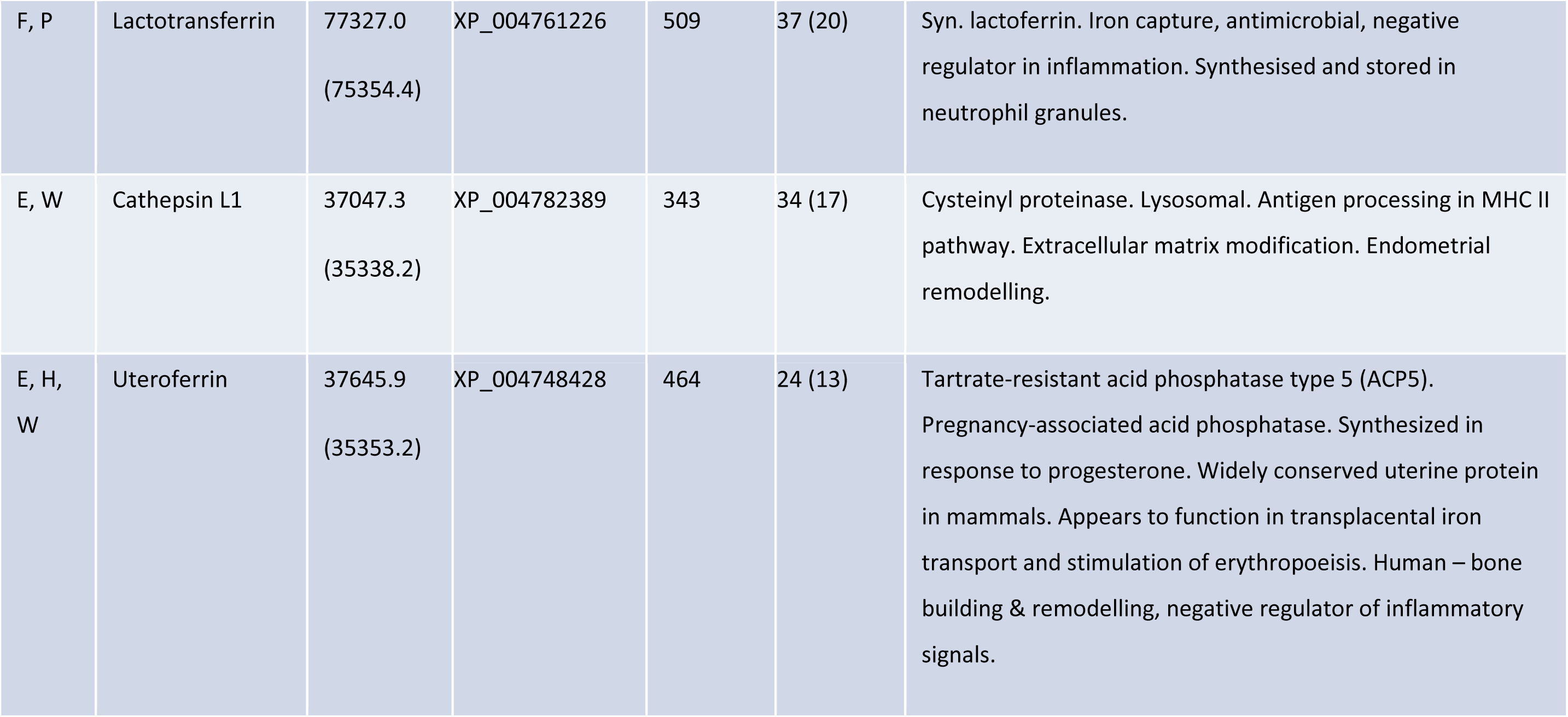
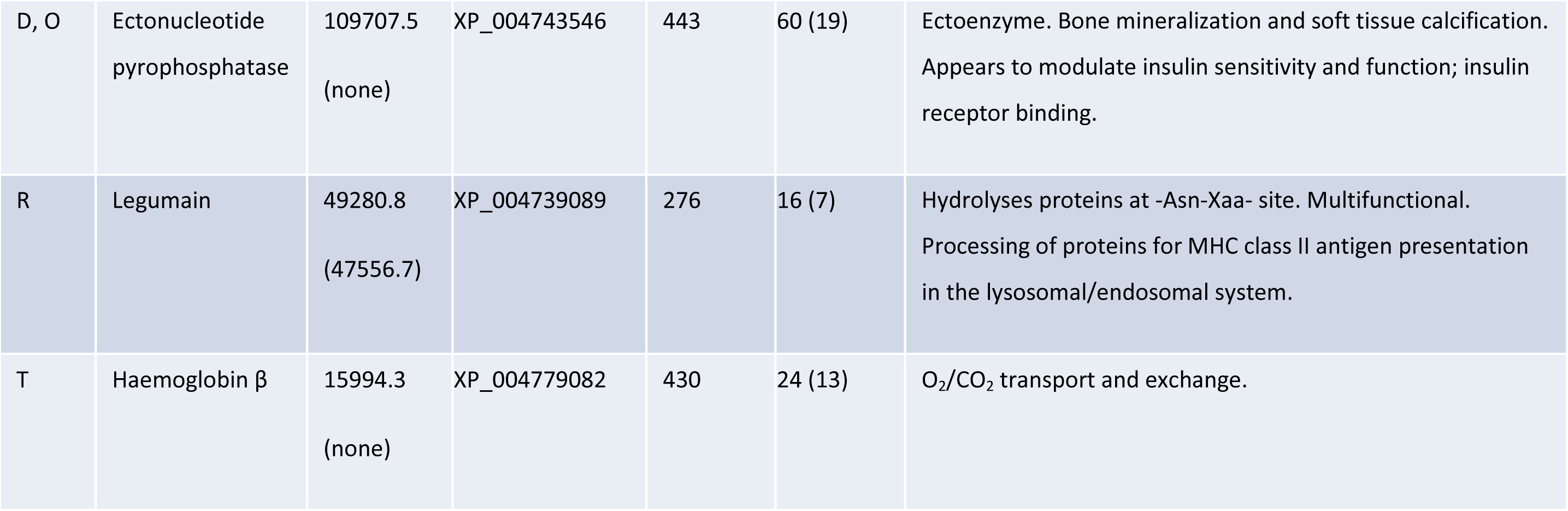
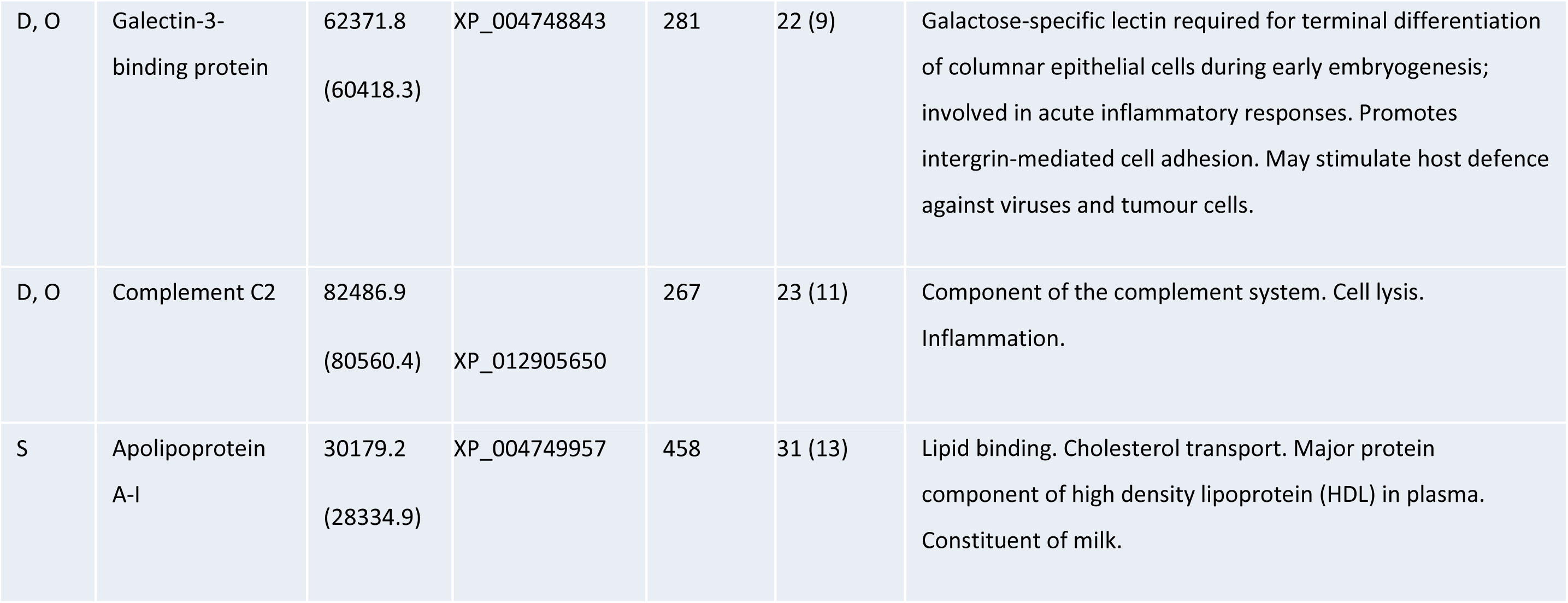
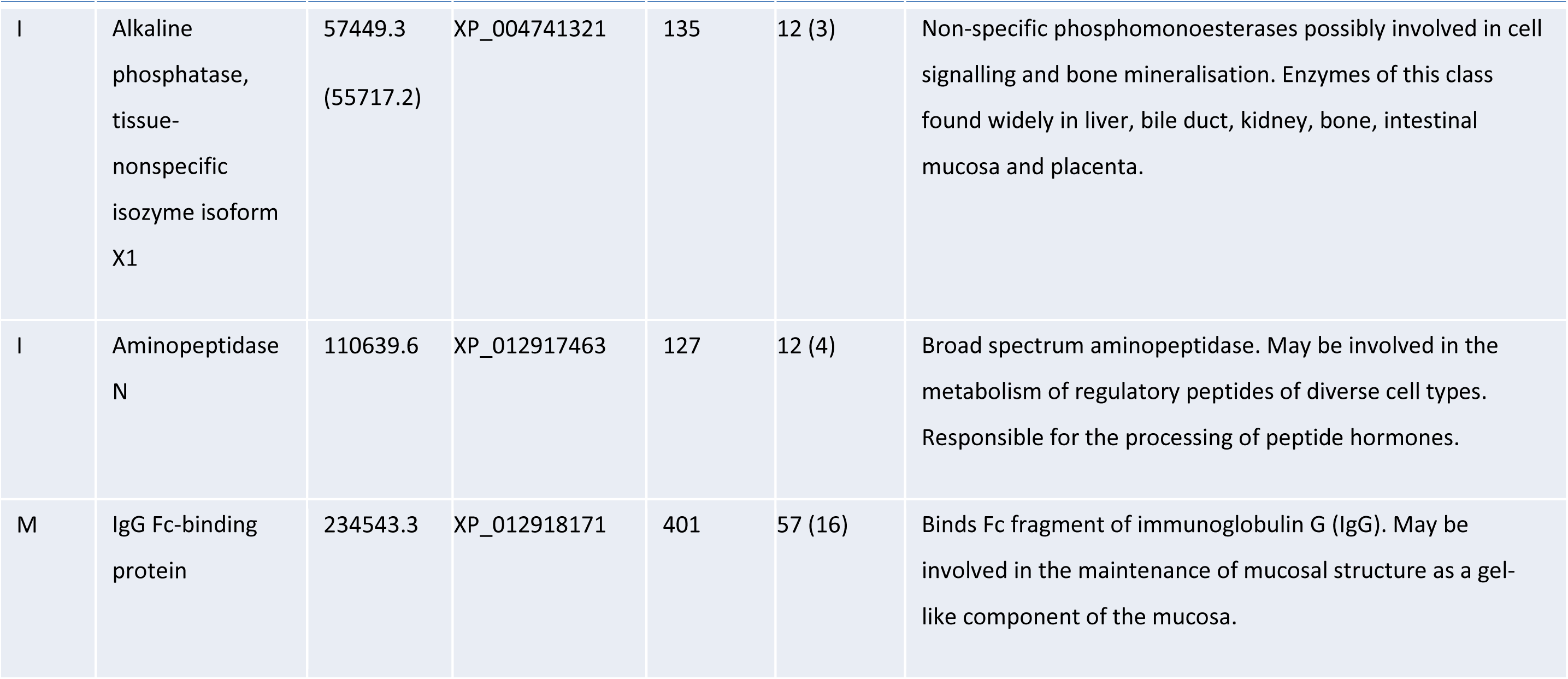
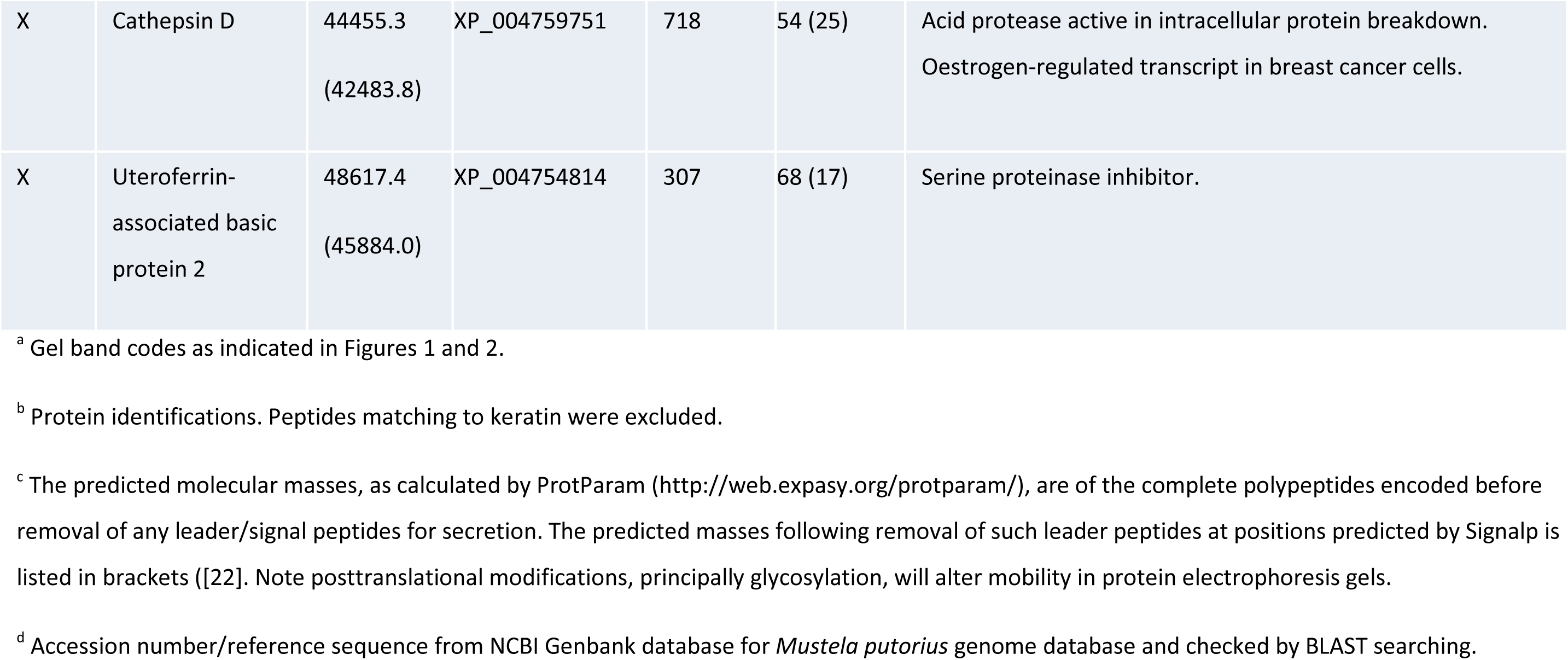
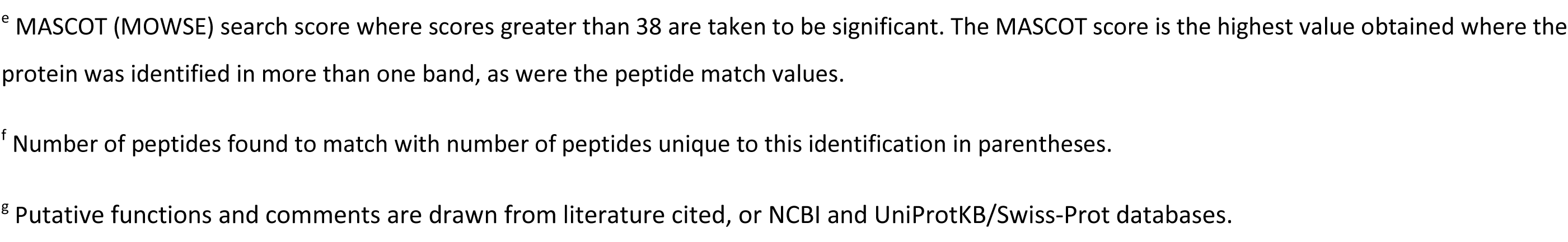
Identification of the proteins isolated from bands excised from the protein electrophoresis gels indicated in Figures 1 and 2.

The two bands with the highest M_r_, bands A and B, were both identified as α-2-macroglobulin (α-2M), a was their reduced form in band J. These sizes are larger than expected from the protein’s polypeptide mass with its predicted secretory signal peptide removed, and may be due to glycosylation and/or covalent linkage with other entities. The other proteins that appeared at high relative amounts around the time of implantation included lipocalin-1 (bands G, U; also known as tear lipocalin, and an enzyme of the ectonucleotide pyrophosphatase family, also represented in bands D, O), cathepsin L1 (bands E, W), uteroferrin (bands E, H, W), and zinc-α-2-glycoprotein (band W). Transferrin (bands F, P), α_1_-antitrypsin (bands C, R, X), and lactoferrin (bands F, P) remained at constant levels relative to serum albumin throughout the sampling period.

## 4. Discussion

The proteins secreted into the uteri of pregnant European polecats changed dramatically in abundance over the approximately 12 days between mating and the time of implantation by the embryo. The repertoire of proteins shows some similarities with that found in other groups of mammals, presumably reflecting a theme inherited from a common ancestor, or convergent evolution in order to satisfy similar functional requirements. In European polecats, however, the appearance of α_2_M was particularly pronounced. Our analysis also identified an abundance of lipocalin-1, which protein has not been reported as a uterine secretion before.

### 4.1 α_2_-macroglobulin

α_2_M was the most notable protein to appear in abundance shortly before the time of implantation. It is closely related to pregnancy zone protein (PZP) and three members of the complement system [23]. It is the largest non-immunoglobulin protein in circulation in mammalian blood, comprises an approximately 160kDa polypeptide that is glycosylated, and usually occurs in circulation as a tetramer comprising a pair of non-covalently associated disulphide-linked dimers [23]. It is synthesised mainly in the liver, but also in the decidua and endometrium in mice and possibly also in humans, and is thought to be important for implantation [12, 24-27].

α_2_M is a minor acute-phase inflammation reactant in several species [28]. It operates via two mechanisms as a broad-spectrum proteinase inhibitor [23]. In one mechanism, an active proteinase cleaves a bait peptide that causes α_2_M to fold around and encapsulate the target enzyme. In the other, the target proteinase is covalently bound via a thioester bond. These mechanisms trap proteinases of all classes but do not necessarily inactivate them, instead removing them from access to intact proteins.

In acute-phase and inflammatory responses, the proteinases targeted by α_2_M are thought to be those of pathogens, but, significantly, also those released by granulocytes and other immune cells in potentially damaging inflammatory reactions [23]. This function could be directly relevant to placentation, given that the sites of placentation in some mammals exhibit local inflammation-like maternal responses [29-31]. The inflammatory response may be amplified by localized apoptosis of epithelial cells, and proteinase inhibition by α_2_M may minimize damage caused by released enzymes [27, 32-37]. α_2_M also binds several hormones, growth factors and cytokines involved in inflammatory responses, and can modify the biological activity of these signalling molecules [19, 23]. In pregnancy, α_2_M is found in uterine secretions of several species, and has been described to control trophoblast positioning and overgrowth in the mouse [7-12, 18, 19].

Several bands in the gels of polecat uterine flushes contained α_2_M, ranging in size from close to that of the monomeic glycoprotein (band J) to higher-order structures (dimer, tetramer; bands A and B). Some that did not conform to simple monomers, dimers or tetramers of the protein may be covalent α_2_M:proteinase complexes that were not dissembled by the reducing agent (Figure 2). The presence of α_2_M in the uterine lumen could be due to increased permeability of the uterine vessels to plasma proteins or tissue disruption. If plasma leakage were to have contributed to the presence of α_2_M, as would also be the case for blood contamination during sampling, then other major plasma proteins, such as immunoglobulins and complement C3, would be present in proportionately high amounts. None of these were observed, although serum albumin was clearly present at all times. While this is the most abundant plasma protein in terms of relative molarity, it is found in uterine secretions of other species in the absence of other plasma proteins [8, 12]. Moreover, α_2_M is known to be synthesised by uterine tissues in humans and mice (see above).

α_2_M and PZP are closely related proteins. The plasma concentration of PZP increases dramatically in the course of human pregnancy. Its functions are not understood [38], although it is known to have proteinase inhibitory activities similar to those of α_2_M [38, 39]. Despite PZP’s documented synthesis in several reproductive tissues [38], and that a gene encoding a PZP homologue has been predicted in the European polecat genome (NCBI accession XP_012906269.1), our analysis identified no protein that might be identified as PZP in the uterine flushes.

### 4.2. Lipocalin-1

The appearance of lipocalin-1 was unexpected, as it has not previously been described as a component of uterine secretions. Moreover, while other lipocalins have been described in the reproductive tissues of other species, only lipocalin-1 was found in this study of polecat uterine secretions. Also known as tear lipocalin and von Ebner’s gland protein, lipocalin-1 is a member of a large family of proteins that operates predominately extracellularly. In addition to its eponymous presence in tears, lipocalin-1 has been found in a wide range of tissues [40], albeit not in uterine secretions.

Most lipocalins bind small hydrophobic ligands such as retinol, fatty acids, odorants, or pheromones, and some are enzymatic [40, 41]. Others are involved in infection defence, such as α_1_-acid glycoprotein (orosomucoid), which is an acute-phase protein in humans that binds a range of small lipophilic molecules and drugs. Lipocalin-2 (neutrophil gelatinase-associated lipocalin) is also an acute-phase protein and binds bacterial siderophores [42, 43]. Lipocalins associated with reproduction include progestogen-associated endometrial protein (glycodelin, pregnancy-associated endometrial α-2-globulin, placental protein 14) secreted by the human endometrium from mid-luteal phase of the menstrual cycle and during the first trimester of pregnancy; a salivary lipocalin found in pigs is secreted into their uteri at the time of implantation [44]; and equine uterocalin is secreted by the endometrium of mares in the pre-placentation period, during which the equine conceptus is confined within a glycoprotein capsule [13, 14]. Despite all these examples of lipocalins found in reproductive tissues in other species, lipocalin-1 was the only one found in our European polecat samples.

The biochemical properties of lipocalin-1 are known only from the human form, which binds small lipids such as fatty acids and sterols [40]. The protein may therefore be involved in delivering or scavenging lipids at a time when there is significant tissue reorganization and potential for cellular disruption at implantation, as is speculated for the role of porcine salivary lipocalin [8]. Lipocalin-1, like lipocalin-2, is known to bind bacterial siderophores, so it may function as part of the innate immune system in an anti-bacterial role [40, 45]. Human lipocalin-1 also has been recorded as having cysteinyl proteinase inhibitory activity [46], and nuclease activity that can operate against viral genomes [47, 48]. The polecat protein may therefore have multiple protective roles in the uterus, as is thought for its function in tears and epithelia in humans [40, 49].

Taken together, these observations suggest that lipocalin-1 may work alongside α_2_M to protect the uterine environment against infections, and its lipid-binding may involve sequestration of toxic lipid peroxidation products created under condition of oxidative stress [50].

### 4.3 Tissue-protective and embryo support proteins

In addition to α-2M and lipocalin-1, we identified several additional proteins that may protect the uterus against infection and tissue damage associated with implantation. These include the iron-binding proteins lactoferrin (lactotransferrin) and transferrin (serotransferrin), both of which were present at fairly constant levels throughout the sampling period. Lactoferrin in particular is associated with antimicrobial activity by sequestering iron in milk, it is also synthesised and stored in neutrophil granules [51], and it directly attacks bacterial membranes and has anti-viral activity [52-56].

α_1_-antitrypsin was also present throughout, and increased in concentration slightly with time. This is an inhibitor of serine proteinases, is a major acute-phase reactant in humans, and is increasingly recognized as an anti-inflammatory mediator [57]. Its primary function appears to be protection against excessive proteolysis of tissues (e.g. in the lower respiratory tract) by neutrophil elastase in inflammation [58, 59]. It has been observed in pregnant uteri of other species, possibly to control proteinases released during tissue remodelling, trophoblast invasion, apoptosis, or inflammatory processes that may accompany placentation [60, 61].

Other proteins increased with time. Cathepsin L1, a cysteinyl proteinase, is usually confined to lysosomes and is involved in MHC class II antigen processing. Its expression level in uterine tissues is progesterone-dependent, and its importance in placentation has been recognized [60, 62, 63]. Other progesterone-dependent proteins that appeared in the polecat uterine fluid included a member of the ectonucleotide pyrophosphatase family (a lysophospholipase that may be involved in the production of pharmacologically-active lipid mediators), and uteroferrin (pregnancy-associated acid phosphatase, which appears to function in transplacental iron transport and stimulation of erythropoeisis) [16, 64-70].

Curiously, we did not find any dedicated nutrient carrier protein similar to equine uterocalin that might act to support a developing conceptus during a pre-placentation period. Equine uterocalin may, however, be a special adaptation in equids to provide resources for a large, capsule-enclosed conceptus during a prolonged pre-placentation period. Nothing like it has been found in non-equids. Mustelid blastocysts do have extended placentation periods, and they can even distend the uterus before they implant, but they have no glycoprotein capsule intervening between their surfaces and the endometrial surface [6]. Moreover, the surface to volume ratios of mustelid conceptuses are substantially smaller than in equids at maximum growth such that they may not need a specialized nutrient carrier. Equine uterocalin and lipocalin-1 bind a similar set of small lipids [15, 40], but the former is atypically enriched in essential amino acids [15], a feature that is not true of lipocalin-1. We did, however, find other lipid carriers that may supply the polecat conceptus with lipids, such as serum albumin and apolipoprotein A-I.

### 4.4. Conclusion

The protein secretory response of the pre-implantation phase of pregnancy that we observed in European polecats appears primarily to prevent infection, facilitate implantation, and protect against release of deleterious cellular components associated with the tissue trauma of implantation. Some of the lipid-binding proteins we found may also be involved in maternal:embryo communication by transporting insoluble signalling molecules such as progesterone, prostaglandins and leukotrienes, or their precursors (as postulated for equine uterocalin; [13, 15]). It would now be interesting to establish whether species of mustelid that exhibit prolonged obligatory embryonic diapause (e.g. stoats, otters, badgers) show a similar protein secretory response when reactivation of blastocysts occurs and implantation is imminent. On the other hand, many of the proteins identified in this study could fulfil the need to protect and nourish a blastocyst that is free in the lumen and devoid of circulatory support and direct immune protection for a considerable time before it implants.

## Acknowledgements

We thank Anitta Helin who took care of the polecats, checked on mating success, and assisted with sample collection, and Suzanne McGill for expert technical assistance with the proteomics analysis. This study was supported by the University of Eastern Finland, Department of Biosciences, and the University of Glasgow. Particular thanks go to Dr Kati Loeffler of the International Fund for Animal Welfare, who critically reviewed the manuscript, found several errors for which the last author was responsible, and contributed a considerable number of excellent suggestions and changes.

## Author contributions

M.W.K. and H.L. conceived, designed, and organised the study. H.L. collected the samples. M.W.K. and R.J.S.B carried out or oversaw the experiments and analysed the data. M.W.K. wrote the paper in consultation with the other authors. All authors saw and approved the final submitted version.

